# GIP receptor agonism improves dyslipidemia and atherosclerosis independently of body weight in obese mice

**DOI:** 10.1101/2022.03.15.484530

**Authors:** Stephan Sachs, Anna Götz, Brian Finan, Richard D. DiMarchi, Yvonne Döring, Christian Weber, Matthias H. Tschöp, Timo D. Müller, Susanna M. Hofmann

**Affiliations:** Institute for Diabetes and Regeneration, Helmholtz Diabetes Center at Helmholtz Zentrum München, German Research Center for Environmental Health (GmbH), 85764 Neuherberg, Germany; Institute for Diabetes and Obesity, Helmholtz Diabetes Center at Helmholtz Centre Munich & Division of Metabolic Diseases, Technische Universität München, Munich, Germany; Technische Universität München, 80333 Munich, Germany; Novo Nordisk Research Center Indianapolis, Indianapolis, Indiana, USA; Department of Chemistry, Indiana University, Bloomington, Indiana. USA; Institute for Cardiovascular Prevention (IPEK), Ludwig-Maximilians-University Munich, Munich, Germany; DZHK (German Centre for Cardiovascular Research), Partner Site Munich Heart Alliance, Munich, Germany; Department of Angiology, Swiss Cardiovascular Center, Inselspital, Bern University Hospital, University of Bern, Bern, Switzerland; Department of Biochemistry, Cardiovascular Research Institute Maastricht (CARIM), Maastricht University, the Netherlands; Munich Cluster for Systems Neurology (SyNergy), Munich, Germany; German Center for Diabetes Research (DZD), 85764 Neuherberg, Germany; Medizinische Klinik und Poliklinik IV, Klinikum der LMU, 80336 München, Germany

## Abstract

Agonism at the receptors for the glucose-dependent insulinotropic polypeptide (GIPR) is a key component of the novel unimolecular co-agonists which are among the most promising drugs in clinical development for the treatment of obesity and diabetes. The therapeutic effect of chronic GIPR agonism to treat dyslipidemia, and thus to reduce the cardiovascular disease risk, has not been explored yet. Herein we found that treatment with a long-acting acylated GIP analog (Acyl-GIP) reduced dyslipidemia and atherogenesis in male LDL receptor knockout mice. Acyl-GIP administration resulted in smaller adipocytes within the inguinal fat depot and RNAseq analysis of the latter revealed that Acyl-GIP may improve dyslipidemia by directly modulating lipid metabolism in this fat depot. This study identified an unanticipated efficacy of chronic GIPR agonist administration to improve dyslipidemia and cardiovascular disease.

---

Alterations in lipid and cholesterol metabolism are major risk factors for the development of cardiovascular diseases (CVD) in patients with obesity and type-2 diabetes (T2D). Albeit best known for its ability to enhance glucose-stimulation of insulin secretion, the glucose-dependent insulinotropic polypeptide (GIP) also stimulates white adipose tissue (WAT) lipid disposal and reduces inflammation in the brain and the white adipose tissue.^1^ Unimolecular co-agonists at the receptors for GIP and the glucagon-like peptide-1 (GLP-1) are among the most promising drugs in clinical development for the treatment of obesity and diabetes^2^. Notably, GLP-1/GIP not only reduces body weight and improves glucose metabolism with greater efficacy relative to GLP-1R agonism in preclinical^3^ and clinical studies^2^, but also outperforms GLP-1R monotherapy in reducing triglyceride and cholesterol levels.^4^ However, the therapeutic effect of GIPR agonism to treat dyslipidemia, and thus reduce the CVD-risk, has not been explored yet. Particularly, it warrants clarification whether GIP may even improve lipid metabolism independent of its ability to improve body weight and glycemia.

Herein we investigated whether a previously published long-acting acylated GIP analog (Acyl-GIP) improves dyslipidemia and atherogenesis in LDL receptor knockout (LDLR−/−) mice. The synthesis, purification, and characterization of fatty-acylated Acyl-GIP was described previously and was used without any further chemical modification or change in formulation.^3^ To assess whether Acyl-GIP affects lipid metabolism and atherosclerotic plaque formation independently of weight loss, we used a dose of Acyl-GIP (10 nmol/kg) that is subthreshold for reducing body weight and for improving glucose metabolism. Consistent with this, body weight (Figure A), body composition (Figure B) and food intake (Figure C) remained similar between vehicle and Acyl-GIP treated LDLR−/− mice. However, 4-week Acyl-GIP treatment remarkably reduced fasting plasma triglycerides and total cholesterol levels in LDLR^−/−^ mice (Figures D-E). This GIP-mediated clearance of plasma lipids was mainly attributable to a decrease of the VLDL and LDL lipoprotein fractions, while HDL levels remained similar to vehicle treated mice (Figure F). Most importantly, GIP treatment was accompanied by reduced atherosclerotic plaque formation within the aortic valve (Figures G-H) and decreased fat streaks along the descending aorta (Figure I). Comprised of anatomically distinct depots, white adipose tissue (WAT) is essential for lipid deposition. Fat accumulation in subcutaneous (sc) fat harbors little to no risk to develop metabolic complications, whereas expansion of visceral (vis) depots predisposes to the metabolic syndrome. We performed RNA sequencing of vis (visWAT) and sc (scWAT) white adipose depots of LDLR−/− mice to explore treatment induced transcriptional changes at study end. Despite higher GIPR expression in visWAT compared to scWAT (Figure J), Acyl-GIP treatment significantly changed the expression of more genes in scWAT compared to visWAT and was associated with smaller adipocytes in scWAT but not visWAT (Figures K-N). In line with GIP-mediated reduction of plasma lipid levels, genes associated with cholesterol and triglyceride metabolism were among the most significantly down regulated pathways in Acyl-GIP-treated scWAT (Figure O). Moreover, Acyl-GIP treatment decreased the expression of genes within the complement and coagulation cascades as well as the fibrinolysis pathway (Figure O). Acyl-GIP induced alterations in lipid metabolism and WAT gene expression were independent of changes in glucose metabolism (Figures P-Q) and, similarly to human trials^5^, we observed that the Acyl-GIP improved dyslipidemia in LDLR−/− mice was independent of changes in insulin metabolism (Figures R-S) potentially indicating a direct effect of Acyl-GIP on adipose tissue metabolism.

Although GLP-1/GIP co-agonists are one of the most promising drugs to treat obesity and diabetes and have been shown to reduce fasting cholesterol and triglycerides in T2D patients, GIP-dependent contributions to metabolic benefits achieved with this combinatorial therapy remain unclear. This study identified an unanticipated efficacy of chronic Acyl-GIP administration to improve dyslipidemia and CVD in a genetic mouse model of atherosclerosis, potentially contributing to the treatment spectrum of clinically advancing novel polypharmacological approaches to treat obesity and T2D. These findings might initiate future studies to explore the potential of GIP mono- or poly-pharmacology to treat disturbances of lipid metabolism, which potentially contributes to reduced cardiovascular mortality.

## Materials and Methods

The data, materials, and methods supporting the findings of this study are available from the corresponding author on request. All mouse procedures were approved by the local Animal Use and Care Committee and the local authorities of Upper Bavaria, Germany in accordance with European and German animal welfare regulations.

### Animals and diet

#### LDLR−/− *mice*

Mice were double-housed and maintained at 22+/−2°C, 55 +/− 10% relative humidity, and a 12-h light/dark cycle with free access to food and water. Mice were randomly assigned to treatment groups matched for body weight and fat mass. All procedures were approved by the local Animal Use and Care Committee and the local authorities of Upper Bavaria, Germany in accordance with European and German animal welfare regulations.

### Compound synthesis

The synthesis, purification, and characterization of the fatty-acylated GIP mono-agonist was described previously and was used without any further chemical modification or change in formulation.^3^

### Rodent pharmacological and metabolism studies

LDLR−/− mice were treated with daily subcutaneous injections (5 μl/g body weight) in the middle of the light phase at the indicated doses with the indicated durations. Body weight and food intake was measured daily. Whole-body composition (fat and lean mass) was measured via nuclear magnetic resonance technology (EchoMRI, Houston, TX, USA). Fasting blood glucose and intraperitoneal glucose tolerance (ipGTT) was determined after a 6h-fast and 20h after the last injection. For ipGTT, 6-h fasted animals were injected intraperitoneally with 2g glucose per kg body weight. Blood glucose was subsequently measured at time points 0, 15, 30, 60, and 120 min using a handheld glucometer (FreeStyle).

### Biochemical analysis

Tail blood prior ipGTT was collected after a 6h fast using EDTA-coated microvette tubes (Sarstedt) and immediately chilled on ice. Mice were euthanized using ketamine (100mg/kg) and xylazine (7mg/kg) injection after a 4h fast and at least 16h after the last vehicle or compound injection. Sac blood was mixed with EDTA and immediately kept on ice. Plasma was separated by centrifugation at 5000 g at 4 °C for 10 min. Plasma levels of insulin (Crystal Chem, IL, USA), cholesterol (Thermo Fisher Scientific, Waltham, MA, USA) and triglycerides (Wako Chemicals, Neuss, Germany) were measured according to the manufacturers’ instructions. For lipoprotein separation, samples were pooled and analyzed in a fast-performance liquid chromatography gel filtration.

### Histology

Atherosclerotic lesion size was assessed by analyzing cryosections of the aortic root by staining for lipid depositions with Oil-Red-O. In brief, hearts with the aortic root were embedded in Tissue-Tek O.C.T. compound (Sakura) for cryosectioning. Oil-Red-O+ atherosclerotic lesions were quantified in 4 μm transverse sections and averages were calculated from 3 sections. The thoraco-abdominal aorta was fixed with 4% paraformaldehyde and opened longitudinally, mounted on glass slides and stained enface with Oil-Red-O. Aortic arches with the main branch points (brachiocephalic artery, left subclavian artery and left common carotid artery) were fixed with 4% paraformaldehyde and embedded in paraffin. Lesion size was quantified after Hematoxylin and Eosin (H&E)-staining of 4 μm transverse sections and averages were calculated from 3-4 sections.

### Statistics

Statistical analyses were performed using GraphPad Prism8. The Kolmogorov-Smirnov test was used to assess for normality of residuals. The unpaired Student two-tailed t-test was used to detect significant differences. A Grubbs test (α < 0.05) was used to detect significant outliers, which were then excluded from subsequent statistical analysis and figure drawing. P < 0.05 was considered statistically significant. All results are mean ± SEM unless otherwise indicated.

### RNA sequencing

Total RNA was extracted from sc and visceral white adipose tissue (scWAT (n=4/treatment) and visWAT (n=5/treatment), respectively) of vehicle and Acyl-GIP vehicle treated LDLR−/− mice (n = 5) using Qiazol according to the manufacturer’s instructions (Qiazol Lysis Reagent, QIAGEN). The quality of the RNA was determined with the Agilent 2100 BioAnalyzer (RNA 6000 Nano Kit, Agilent). All samples with a RNA integrity number (RIN) had a value greater than 7. For library preparation, 1 μg of total RNA per sample was used. RNA molecules were poly(A) selected, fragmented, and reverse transcribed with the Elute, Prime, Fragment Mix (EPF, Illumina). End repair, A-tailing, adaptor ligation, and library enrichment were performed as described in the TruSeq Stranded mRNA Sample Preparation Guide (Illumina). RNA libraries were assessed for quality and quantity with the Agilent 2100 BioAnalyzer and the Quant-iT Pico- Green dsDNA Assay Kit (Life Technologies). Strand-specific RNA libraries were sequenced as 150 bp paired-end runs on an Illumina HiSeq4000 platform. The STAR aligner* (v 2.4.2a)57 with modified parameter settings (–twopassMode = Basic) was used for split-read alignment against the mouse genome assembly mm10 (GRCm38) and UCSC known Gene annotation. To quantify the number of reads mapping to annotated genes we used HTseq-count (v0.6.0). For differentially testing we followed guidelines reported by Law et al. (Law et al. 2016 10.12688/f1000research.9005.2). Briefly, we excluded genes with zero counts in all samples and further removed genes with a cumulative counts per million in less than five samples. We used the edgeR package for data pre-processing, followed by the limma package with its voom method, linear modelling and empirical Bayes moderation to assess differential expression. We used EnrichR web interface for gene and pathway enrichment. As input, genes with a p-value < 0.05 and a logFC > 0.75 were used.

## Acknowledgments

We thank Luisa Müller, Laura Sehrer, Emiljia Malogajski, Cynthia Striese and Sebastian Cucuruz from the Helmholtz Diabetes Center in Munich and Yvonne Jansen from the institute for Cardiovascular Prevention (IPEK), Ludwig-Maximilians-University Munich, for excellent assistance with in vivo mouse experiments as well as Cynthia Striese for her excellent help preparing and analyzing heart tissues. C.W. is van der Laar-Professor of Atherosclerosis.

---

**Figure:**
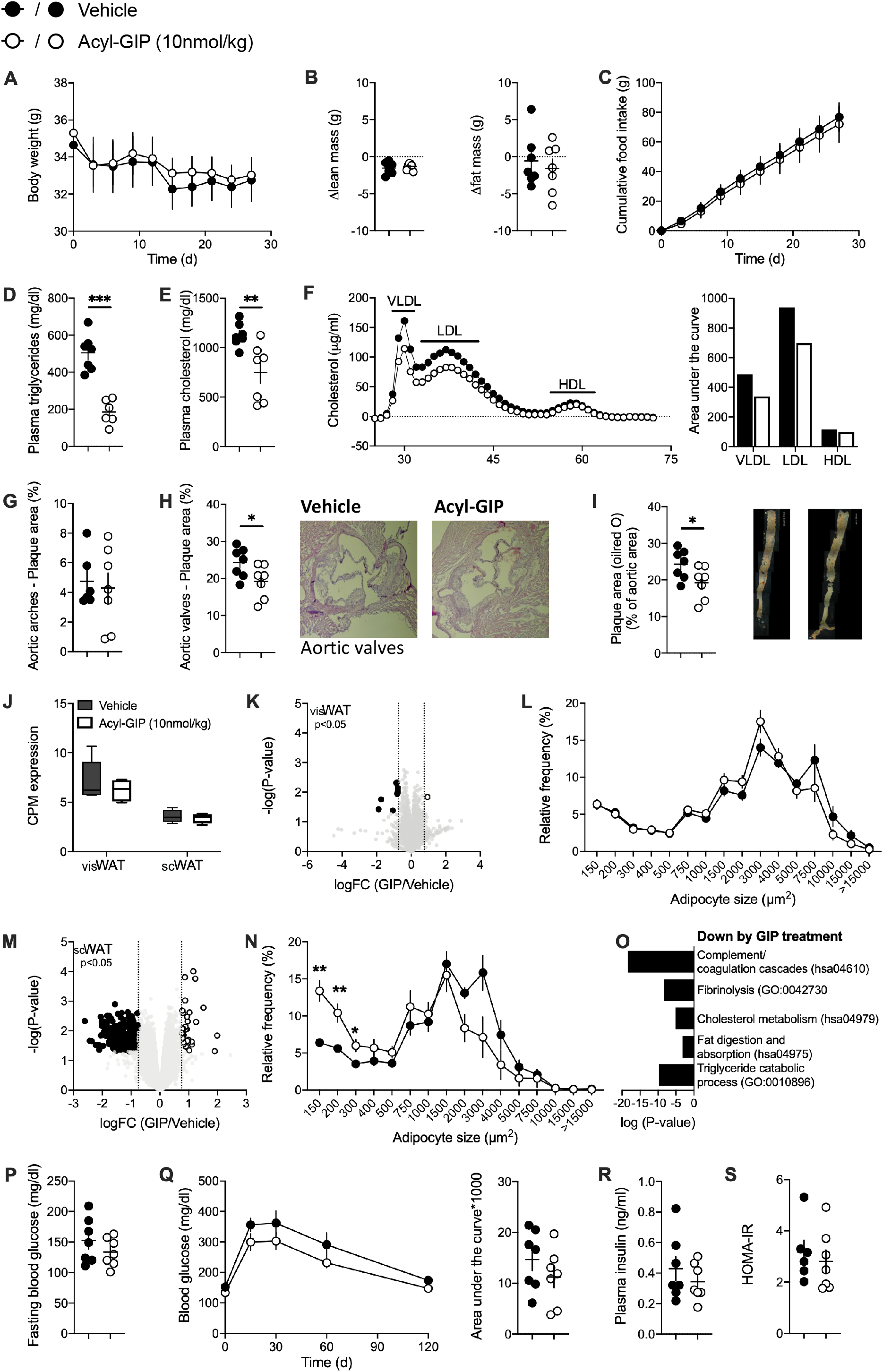
Acyl-GIP ameliorates dyslipidemia in LDLR−/− mice. 8-week old male LDLR−/− mice were fed a western diet high in calories and cholesterol (21% fat, 0.2% cholesterol) for 8 weeks to induce atherogenic dyslipidemia prior treatment start and were maintained on this diet during either daily subcutaneous vehicle or Acyl-GIP injections. (A) Body weight, (B) change of body composition, (C) cumulative food intake, plasma (D) triglycerides, (E) cholesterol and (F) lipoprotein fractions as well as the percentage of plaque area in aortic arches and valves and along the descending aorta of male LDLR−/− mice treated daily with either vehicle or Acyl-GIP via subcutaneous injections for 28 days. n = 7. (J) Relative visWAT and scWAT GIPR gene expression of vehicle (visWAT n=5; scWAT n=4) and Acyl-GIP treated LDLR−/− mice (visWAT n=5; scWAT n=4). (K) Volcano plot showing differential expression and its significance (-log10(p-Value), limma-trend) and (L) frequency distribution of adipocyte cell sizes (μm2) of visWAT from Acyl-GIP (RNA sequencing n=5; histology n=7) versus vehicle (RNA sequencing n=5; histology n=6) treated LDLR−/− mice. (M) Volcano plot showing differential expression and its significance (-log10(p-Value), limma-trend) and (N) frequency distribution of adipocyte cell sizes (μm2) of scWAT from Acyl-GIP (RNA sequencing n=5; histology n=4) versus vehicle treated LDLR−/− mice (RNA sequencing n=5; histology n=4). (O) Selected gene ontologies (p<0.0001) and KEGG pathways (P value < 0.001) that are down regulated by Acyl-GIP in scWAT. (P) Fasting blood glucose, (Q) interperitoneal glucose tolerance test (ipGTT), (R) fasting insulin levels and (S) the homeostatic model assessment of insulin resistance (HOMA-IR) of male LDLR−/− mice treated daily with either vehicle or Acyl-GIP via subcutaneous injections for 28 days. n = 7. Except for fasting glucose and ipGTT (performed at study day 22), plasma parameters were measured from sac blood at study end (day 28). n = 7. Data represent means ± SEM. *P < 0.05, **P < 0.01, *** P < 0.001, determined by unpaired two-sided t-test.

